# Influence of Macrocyclization Strategies on DNA-Encoded Cyclic Peptide Libraries

**DOI:** 10.1101/2025.04.10.648295

**Authors:** Hanqing Zhao, Xue Li, Xueyu Yao, Shijie Zhang, Yandan Bao, Weiwei Lu, Minyan Xing, Xudong Wang, Xuan Wang, Yujun Zhao, Qian Chu, Xiaojie Lu

## Abstract

Discovery of cyclic peptide hits using DNA-encoded libraries (DELs) has recently been extensively researched, with significant efforts directed toward developing DEL-compatible macrocyclization methods. To investigate how different cyclic linkers influence DEL selection outcomes, we constructed eight distinct sub-libraries and screened them against two protein targets, MDM2 and GIT1, resulting in two representative yet contrasting scenarios. Validation studies for MDM2 revealed that structural similarity patterns observed across multiple sub-libraries could enhance confidence in the authenticity of identified hits. Notably, **cmp-10** demonstrated potent inhibitory activity, exhibiting an inhibition constant of 11 nM. In contrast, selections against GIT1 produced discrete enrichment patterns. From these outcomes, two exemplary compounds were selected and validated through both on-DNA and off-DNA modes. **cmp-17** was confirmed to bind specifically to the desired binding pocket, displaying a dissociation constant of 1.22 μM. Furthermore, ITC experiments using mutant GIT1 proteins (GIT1^L271A/L279A^ and GIT1^L271L/L279L^) provided additional insights into the mechanism by which **cmp-17** disrupts the interaction between GIT1 and β-Pix.

## INTRODUCTION

In recent years, cyclic peptide therapeutics have garnered significant attention, prompting researchers to extensively explore and expand their applications across various fields including orally administered drugs, peptide-drug conjugate, diagnostic probes, and so on^1-3^. Despite this growing interest, the efficient discovery of cyclic peptide candidates remains a considerable challenge.

Currently, most clinically approved peptide drugs are derived from natural products or their derivatives. However, relying solely on this approach may not sufficiently meet the increasing demands of pharmaceutical development. Consequently, researchers have focused on developing and applying novel technologies, ranging from entirely rational design to serendipitous discovery methods. Among these, genetically encoded display technologies, such as phage display and mRNA display, have emerged as particularly powerful tools for identifying *de novo* peptide candidates. ^4-5^ Both methods excel at rapid generation and screening of extensive peptide libraries against diverse molecular targets, leading to the discovery of novel peptide candidates. Several peptides identified through these technologies have already been approved by the FDA or are currently in clinical trials. ^6-11^ Phage display utilizes engineered bacteriophages to express randomized peptide libraries on their surface proteins, whereas mRNA display employs *in vitro* translation systems, linking genotype to phenotype by covalently attaching nascent peptides to their encoding mRNA *via* puromycin. Both technologies traditionally produce peptides composed of the 20 natural amino acids. However, natural peptides often suffer from metabolic instability and poor membrane permeability. To overcome these limitations, incorporation of noncanonical amino acids^12^ and more rigid cyclic scaffolds has significantly improved peptide properties. For instance, phage display has been combined with genetic code expansion techniques to incorporate noncanonical amino acids^13^, while mRNA display employs flexizymes to enable similar incorporation^14-15^. Despite these advances, the high technical complexity of these methods limits their broader adoption, particularly among new users.

In contrast, chemically synthesized libraries readily accommodate diverse non-canonical amino acids and enable various macrocyclization methods for cyclic peptides synthesis. By integrating DNA barcodes with synthetic molecules, DNA-encoded library technology (DELT) allows rapid generation of compound libraries suitable for subsequent selection campaigns. DELT has thus attracted considerable attentions, emerging as an essential complement to display technologies and greatly enhancing the discovery of novel peptide candidates with improved pharmacokinetic profiles. ^16^

Historically, DELT was primarily applied to small molecule discovery, with several identified candidates progressing into clinical trials. Following validation of its feasibility and reliability, DELT rapidly gained widespread adoption in both academic and industrial research groups. ^17-19^ Prior to 2020, numerous efforts were already underway to identify cyclic peptide binders using DELT. A notable example is DNA-templated synthesis (DTS), pioneered by Liu’s group, which demonstrated distinctive capability for constructing cyclic peptide libraries. ^20^ Building on early successes, Ensemble Therapeutics, co-founded by Professor Liu, continued to advance cyclic peptide development. However, with further technological advancements, DNA-recorded synthesis gradually became the predominant method for generating DNA-encoded libraries, leading to several exemplary DELs. In 2018, GSK reported the first DNA-recorded cyclic peptide libraries, utilizing six synthetic cycles to produce up to 2.4 trillion uniquely encoded compounds. ^21^ Concurrently, the Neri/Scheuermann group developed an alternative strategy involving the initial attachment of pre-cyclic scaffolds to DNA tags, followed by site-specific chemical modifications. ^22^ Additionally, solid-phase DEL methods were developed and optimized by Paegel^23^, Kodadek^24^, and Lim^25^ research groups. This approach provided significant advantages, including compatibility with water-sensitive reactions and the capability for functional screening.

Since 2020, growing enthusiasm for cyclic peptide therapeutics has accelerated further exploration of DEL technology. Consequently, numerous DEL-compatible methodologies, encoded libraries, and cyclic peptide hits exhibiting exceptional inhibitory activities or binding affinities have been reported. However, a cursory review of recent DNA-encoded chemical library (DECL) literature reveals an intensive focus on developing on-DNA macrocyclization methods^26-33^, with relatively few studies specifically addressing data analysis for cyclic peptide DELs^34^. Traditionally, DEL selection analysis relies on counting the frequency of enriched features and visualizing the results in a three-dimensional scatter plot. By deconvoluting the structure-enrichment relationship—where the contribution of individual building blocks is identified as singletons, lines, or planes—potential hit compounds can be analyzed *in silico*, selected, and subsequently resynthesized off-DNA for validation. Several data-driven approaches have also been developed to manage small-molecule DEL data outputs, including normalized Z-scores^35^, Poisson distribution normalization^36^, “active-fraction” calculations^37^, and computational tools such as DELdenoiser^38^. However, the relatively flexible conformations and hydrophobic characteristics of cyclic peptides pose unique analytical challenges in reliably identifying genuine binders while minimizing interference from DNA tags.

This paper presents two case studies involving the selection of cyclic peptide binders from DNA-encoded cyclic peptide libraries (DECPLs) targeting MDM2 and GIT1 proteins. MDM2 (murine double minute 2) ^39^ is a key negative regulator of the tumor suppressor p53 by promoting its ubiquitination and subsequent degradation. MDM2 inhibition has been extensively investigated as potential anti-cancer therapy, with several inhibitors demonstrating promising results in both preclinical and clinical studies. Given the abundance of available structural and biochemical data, MDM2 is commonly employed as a model protein to evaluate novel screening technologies. ^40^ GIT proteins, including GIT1 and GIT2, function as GTPase-activating proteins (GAPs) for Arf family GTPases, while their binding partners, PIX proteins (α-PIX and β-PIX) act as guanine nucleotide exchange factors (GEFs) for Rho family GTPases. ^41^ Extensive evidence indicates that the GIT1/β-PIX interaction strongly correlates with cancer progression and metastasis across various human malignancies. Structural analysis of the publicly available crystal structure of the GIT/PIX complex reveals an extremely shallow binding interface, making modulation of this protein-protein interaction (PPI) particularly challenging. Cyclic peptides offer a promising approach to address such difficult PPIs. Thus, we aimed to screen the GIT1 protein using DNA-encoded cyclic peptide libraries.

The DNA-encoded cyclic peptide libraries (DECPLs) employed in this study consist of eight sub-libraries. These sub-libraries share the same building blocks (BBs) but feature different cyclization strategies, either through oxidative disulfide bond formation or thiol-electrophile cross-linking with various bifunctional linkers. Our selection results for MDM2 and GIT1 exemplify two typical scenarios encountered in DEL screening. Repeatedly enriched building block (BB) combinations across multiple sub-libraries can enhance confidence in the authenticity of identified hits. Conversely, isolated enrichments appearing in individual sub-libraries require rigorous validation to confirm the reliability of the selected compounds.

## RESULTS AND DISCUSSION

### The generation of DNA-Encoded Cyclic peptide Libraries (DECPLs) and the affinity selection workflow

The DNA-encoded cyclic peptide libraries (DECPLs) were generated following the synthetic workflow illustrated in **Figure 2a**. Library synthesis began with a bifunctional “headpiece,” followed by iterative split-and-pool combinatorial cycles involving on-DNA chemical reactions and enzymatic ligation. For each integrated DNA-tagged library member, DNA tags were elongated from short DNA sequences catalyzed by T4 ligase. Concurrently, cyclic peptide moieties were assembled using amino acid building blocks (BBs) *via* DNA-compatible amidation and subsequent deprotection, followed by final macrocyclization. The resulting cyclic peptide moieties used in this study featured three structural elements: (1) Three variable positions, each randomly incorporating one amino acid building block selected from a repertoire of 536 amino acids, ultimately generating a cyclic peptide library containing approximately 154 million unique members; (2) Two thiol-containing units, cysteine and 3-(pyridin-2-yldisulfanyl)propanoic acid, positioned at the N- and C-termini, , respectively; (3) Bis-electrophilic linkers used to react with the thiol groups, forming thioether bridges in libraries **DEL-1** ∼ **DEL-7**, or alternatively, creating the disulfide-cyclized peptide library in **DEL-8** *via* oxidative cyclization.

**Figure 1.**
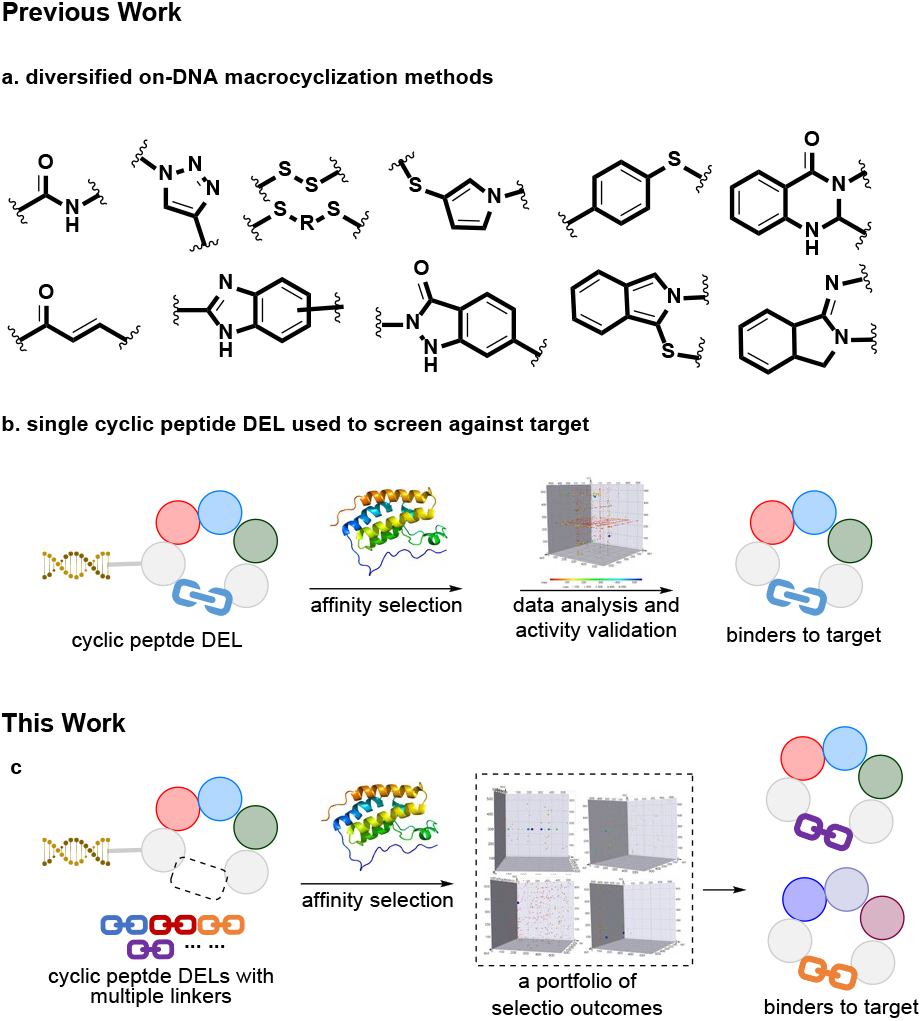
(a) exemplary DNA-compatible reactions developed in recent years. (b) affinity binders selected from individual enrichment data. (c) affinity binders selected from collected enrichment data.

**Figure 2.**
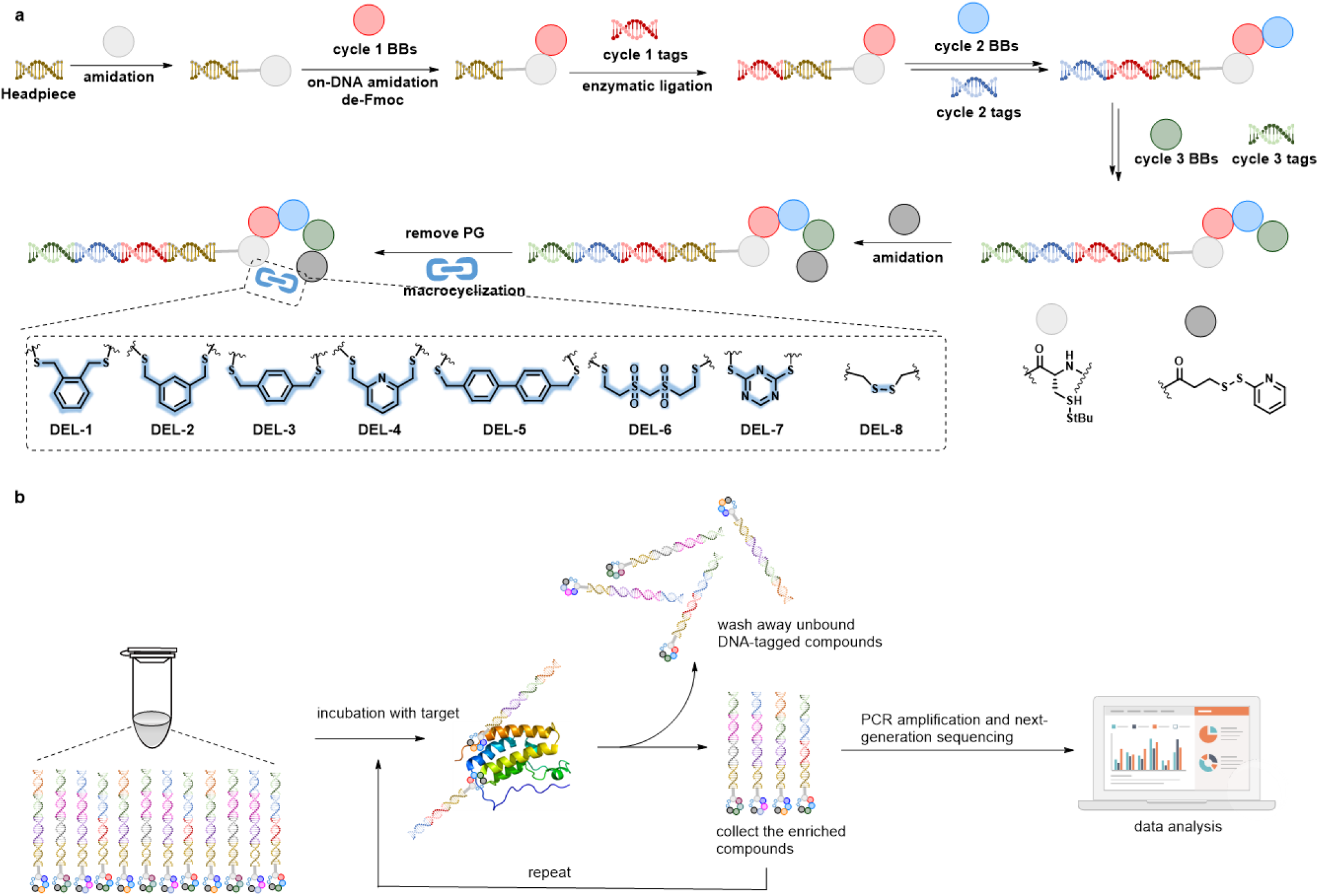
(a) the synthetic scheme of DNA-encoded cyclic peptide libraries. (b) the routine workflow of affinity selection campaign.

The typical affinity selection process is illustrated in **Figure 2b**. In the case of protein targets containing a His-tag, the proteins were immobilized onto HisPur™ Ni-NTA Magnetic Beads and then incubated with the pooled libraries **DEL-1** ∼ **DEL-8**. Non-binding library members were removed by washing, and bound peptides were subsequently eluted by heating. The eluted binders were collected and subjected to the next selection round. This affinity selection cycle was repeated three times. After the final round, the DNA tags linked to the enriched binders were amplified *via* PCR and identified by next-generation sequencing (NGS). The resulting NGS data were decoded, analyzed in silico, filtered using informatics-based cutoff criteria, and visualized as three-dimensional scatter plots for further analysis.

### Affinity Selection of DECPLs and binding validation for MDM2 binders

The selection outcome of DECPLs against the MDM2 target is illustrated in **Figure 3**, with **DEL-1** serving as an example of the analysis process. In the three-dimensional scatter plot (**Figure 3a**), each axis represents a building block (BB1, BB2, and BB3) corresponding to reaction cycles 1, 2, and 3, respectively. Each dot represents an individual encoded compound, and the size of each dot positively correlates with the “select_fold_to_mean” value of the compound, defined as the ratio of the enrichment value of each library member to the mean enrichment value within the selection group. This normalization parameter was chosen to facilitate the comparison of selection outcomes across different DEL libraries. To maximize the probability of identifying authentic hit compounds, we aimed to triage candidates based on quantitative readouts, including sequence counts and structural similarity among enriched features. The fundamental principle underlying DEL selection data analysis is that molecules with higher affinity to the target protein become preferentially enriched, resulting in higher enrichment values. In the plot, several peptide hits lined up as highlighted in the red box, indicating that the encoded compounds sharing common building blocks at cycle 2 (blue) and cycle 3 (green). Analysis of the cycle 1 building blocks within this cluster revealed a structural motif characterized by a β-phenyl moiety and its derivatives. Such structural similarity patterns among enriched binders enhance confidence in the validity of the identified molecules. Consequently, candidate molecules for further validation were selected from this enriched cluster of chemically related compounds. For example, **cmp-01** (BB combination 231_246_92) was selected, resynthesized off-DNA, and demonstrated robust binding with MDM2 protein with a Ki value of 72 nM in a competitive binding assay.

**Figure 3.**
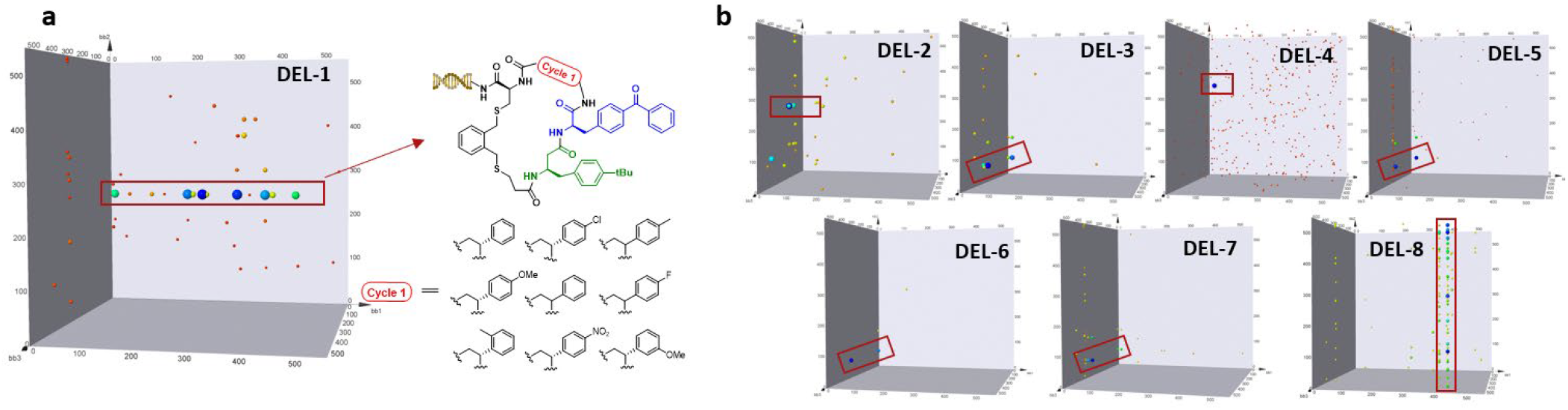
The visualized 3D plots of MDM2’s selection outcomes.

Notably, the other seven libraries (**DEL-2** ∼ **DEL-8**) revealed completely different structural enrichment patterns compared to **DEL-1** (**Figure 3b**). In **DEL-2**, the highlighted encoded compounds shared the same building blocks at cycle 1 (BB No. 62) and cycle 2 (BB No. 273). The corresponding highly enriched molecule, **cmp-03**, was selected for off-DNA resynthesis and subsequent validation, exhibiting a K_i_ value of 2.9 μM (**Table 1**). Conversely, **DEL-4** illustrated a contrasting scenario, where only a single specific BB combination showed a high enrichment ratio. Such singleton hits may represent genuine binders in the context of the entire BB repertoire. The corresponding compound **cmp-06** was validated and demonstrated a K_i_ value of 0.363 μM. Although **DEL-8** exhibited a clear structural enrichment pattern, no compounds were chosen for validation due to the presence of an unfavorable trityl (Trt) moiety in the identified BBs, thus lowering the synthetic priority for this series of structurally similar binders.

**Table 1.**
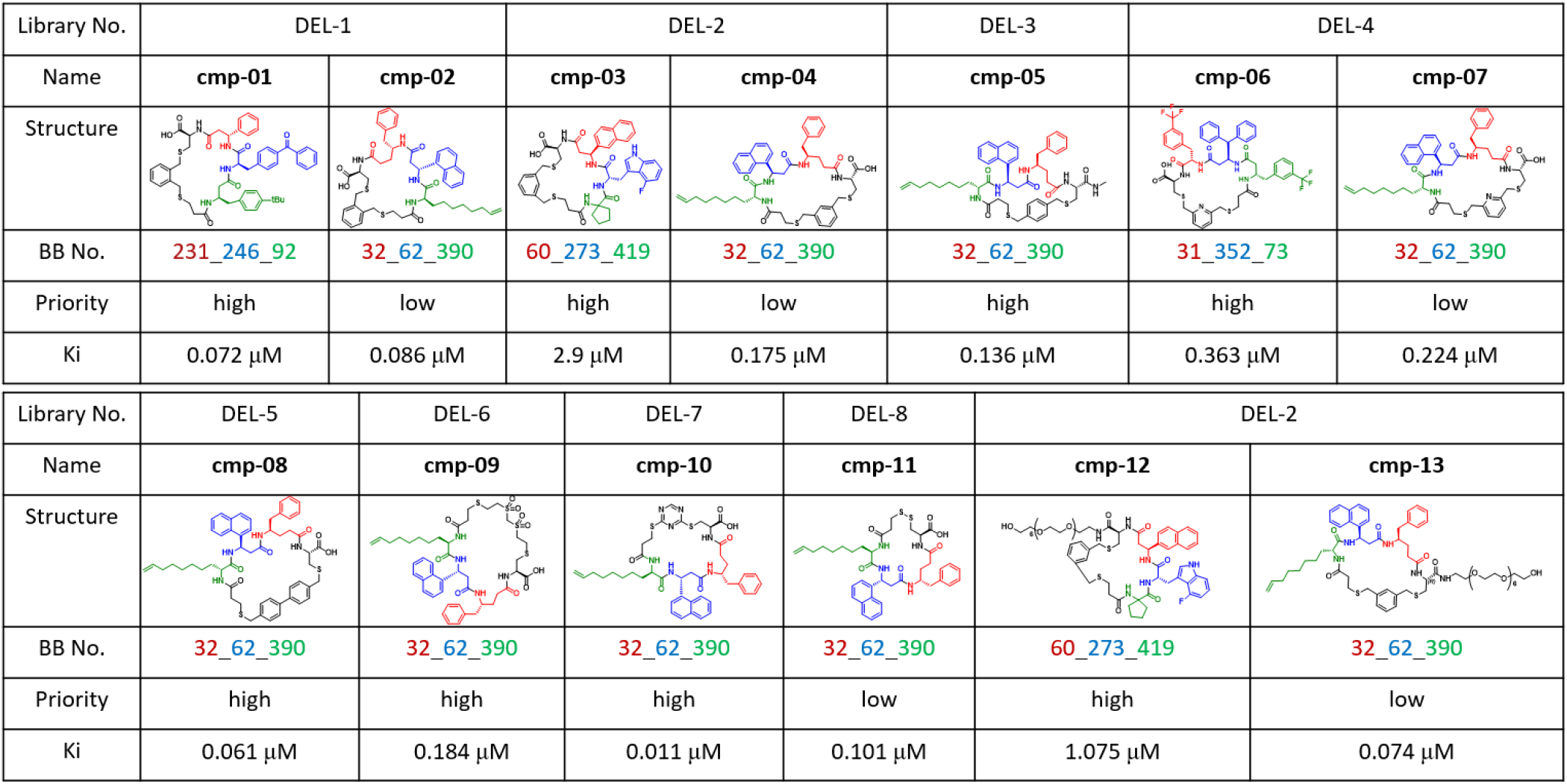
Validation of the selected compounds by off-DNA synthesis and FP assay.

| Library No. | DEL-1 |  | DEL-2 |  | DEL-3 | DEL-4 |  |
| --- | --- | --- | --- | --- | --- | --- | --- |
| Name | cmp-01 | cmp-02 | cmp-03 | cmp-04 | cmp-05 | cmp-06 | cmp-07 |
| Structure |  |  |  |  |  |  |  |
| BB No. | 231_246_92 | 32_62_390 | 60_273_419 | 32_62_390 | 32_62_390 | 31_352_73 | 32_62_390 |
| Priority | high | low | high | low | high | high | low |
| Ki | 0.072 μM | 0.086 μM | 2.9 μM | 0.175 μM | 0.136 μM | 0.363 μM | 0.224 μM |

| Library No. | DEL-5 | DEL-6 | DEL-7 | DEL-8 | DEL-2 |  |
| --- | --- | --- | --- | --- | --- | --- |
| Name | cmp-08 | cmp-09 | cmp-10 | cmp-11 | cmp-12 | cmp-13 |
| Structure |  |  |  |  |  |  |
| BB No. | 32_62_390 | 32_62_390 | 32_62_390 | 32_62_390 | 60_273_419 | 32_62_390 |
| Priority | high | high | high | low | high | low |
| Ki | 0.061 μM | 0.184 μM | 0.011 μM | 0.101 μM | 1.075 μM | 0.074 μM |

**DEL-3, DEL-5, DEL-6**, and **DEL-7** demonstrated comparable selection outcomes, with the BB combination 32_62_390 consistently producing high enrichment ratios. All corresponding compounds (**cmp-05, cmp-08, cmp-09**, and **cmp-10**) exhibited nanomolar binding affinities, among which **cmp-10** showed the highest affinity (K_i_ = 11 nM). Interestingly, although the BB combination 32_62_390 showed relatively low enrichment ratios in **DEL-1, DEL-2, DEL-4**, and **DEL-8**, we resynthesized the corresponding cyclic peptides (**cmp-02, cmp-04, cmp-07**, and **cmp-11**) to compare their binding affinities with highly enriched compounds. Surprisingly, both **cmp-02** and **cmp-07** displayed good binding affinities comparable to **cmp-01** and **cmp-06**, respectively. Moreover, **cmp-04** exhibited even better binding affinity compared to **cmp-03**, contrary to the initial selection outcomes. To further validate these results, we synthesized PEG-modified compounds (**cmp-12** and **cmp-13**) to mimic the spatial context of DNA-tagged compounds. Although both PEG-modified analogs showed improved binding affinities, **cmp-13** still exhibited superior affinity compared to **cmp-12**. These findings indicate that the correlation between the selection enrichment of DNA-tagged compounds and the binding affinities of off-DNA compounds is more complicated than initially acknowledged. Numerous factors complicate this relationship, including hydrophilicity, steric hindrance, and altered binding conformations induced by the DNA tags. Such complexity is particularly pronounced in cyclic peptide DELs compared to small-molecule DELs due to their inherently more flexible binding conformations.

These results also challenged our previous protocol for binder selection. If we had confined our screening to DEL-2 alone, the BB combination 32_62_390 would have been overlooked for subsequent validation due to its low enrichment ratio. This highlights the importance of considering structural similarity patterns across multiple hit series or different libraries, as such cross-library comparisons can increase confidence in the authenticity of observed hits.

### Affinity Selection and validation for GIT1 binders

The GIT1 target, screened using the same libraries (**DEL-1** ∼ **DEL-8**), presented a contrasting scenario compared to the MDM2 selection results, with no consistent enrichment patterns observed across libraries (**Figure 4a**). Thus, cross-referencing between libraries was not feasible to assist in identifying authentic hit compounds. This phenomenon is not unique and has been observed with other targets as well. Consequently, a standardized protocol was employed to select potential binders for further validation.

**Figure 4.**
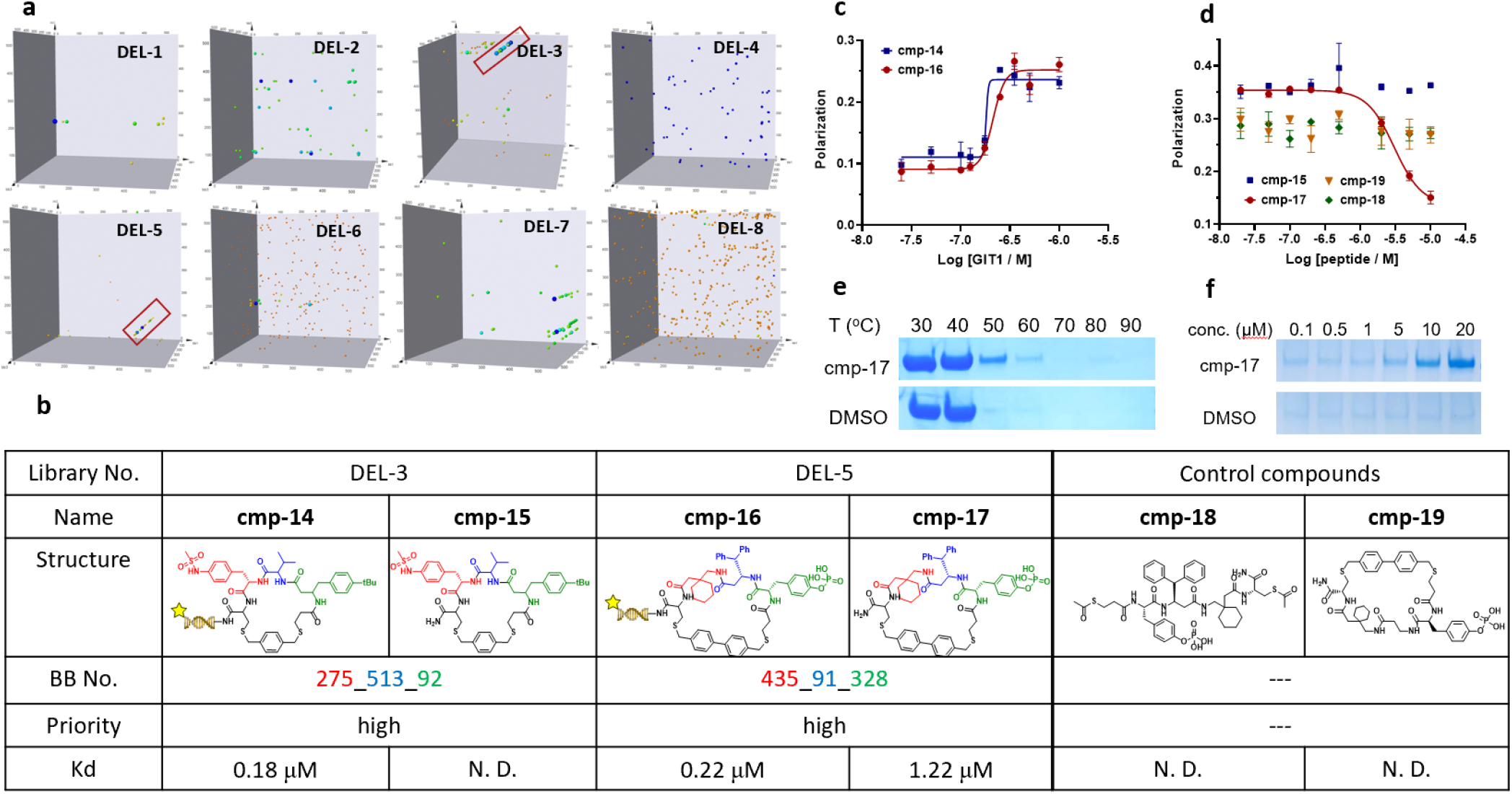
(a) the visualized 3D plots of GIT1’s selection outcomes. (b) validation results of **cmp-14** ∼ **cmp-19** through both on-DNA and off-DNA modes. (c) the binding plot of DNA-tagged compounds. (d) the binding plot of off-DNA compounds. (e) the TSA assay result of **cmp-17**. (f) the dose-dependent assay results of **cmp-17**.

As illustrated in **Figure 4a, DEL-3** and **DEL-5** displayed distinct enriched clusters (highlighted in red). The corresponding BB combinations, 275_513_92 from **DEL-3** and 435_91_328 from **DEL-5**, were selected for validation (**Figure 4b**). We synthesized both FITC-labeled DNA conjugates (**cmp-14** and **cmp-16**) and their corresponding non-labeled off-DNA peptides (**cmp-15** and **cmp-17**). Fluorescence polarization (FP) binding assay revealed that both **cmp-14** and **cmp-16** exhibited robust binding towards the GIT1 protein, with K_d_ values of approximately 181 nM and 212 nM, respectively (**Figure 4c**). To further evaluate the cyclic peptide hits and rule out the potential interference from the DNA tag, we conducted a competitive FP experiment using the non-labeled peptides (**cmp-15** and **cmp-17**) to compete for GIT1 binding with the fluorescein-labeled β-Pix peptide (**Figure 4d**). Notably, **cmp-17** demonstrated effective competition against the β-Pix peptide with a K_i_ value of 1.22 μM, confirming its ability to specifically target and disrupt the GIT1-β-Pix interaction. However, **cmp-15** failed to displace the β-Pix peptide, presumably because the peptide binds at a distinct site distal from GIT1-β-Pix interaction or the fused DNA sequence in **cmp-14** may contribute to the binding process. Therefore, we focused on **cmp-17** from **DEL-5** for subsequent investigations.

First, we employed a thermal shift assay (TSA) to further assess the binding of **cmp-17** to GIT1. Temperature screening revealed that **cmp-17** significantly enhanced the thermal stability of recombinant GIT1 between 50 °C and 60 °C (**Figure 4e**). In addition, **cmp-17** exhibited dose-dependent stabilization of GIT1 at the low micromolar range, further confirming its interaction with the target protein (**Figure 4f**). Next, we designed and synthesized two negative control peptides to dissect the structural determinants of binding. **Cmp-18** is a linearized version of **cmp-17**, with both thiol groups methylated to prevent cyclization. Meanwhile, **cmp-19** is a cyclic peptide derived from **cmp-17** by depleting the benzhydryl side chain, which is presumably a key binding residue (**Figure 4b**). In the competitive FP assay, neither control peptide exhibited detectable activity (**Figure 4d**). Furthermore, isothermal titration calorimetry (ITC) experiments confirmed no measurable binding between GIT1 and either **cmp-18** or **cmp-19**, whereas **cmp-17** showed a dissociation constant K_d_ of 1.28 μM, highlighting its binding ability and specificity for GIT1 (**Figure 5**).

**Figure 5.**
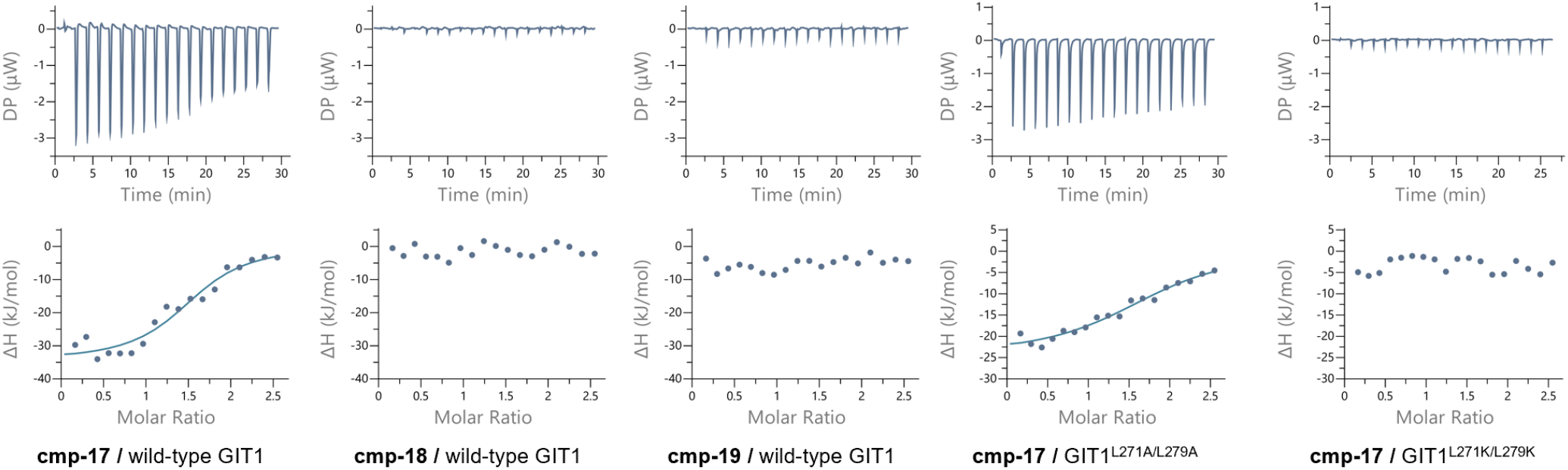
the ITC assay results of the off-DNA compounds.

To further characterize how **cmp-17** disrupts the GIT1-β-Pix interaction, we aimed to identify the key residues in GIT1 that mediate binding. Competitive FP assay indicated that **cmp-17** binds at the same site as the β-Pix peptide. According to the crystal structure of the GIT1-β-Pix complex, two conserved leucine residues (Leu271 and Leu279) form a hydrophobic interface and are essential for complex assembly. Therefore, we mutated these two residues to alanines (GIT1^L271A/L279A^) to reduce the hydrophobicity at the binding interface. In addition, GIT1^L271K/L279K^ was also obtained by introducing lysine substitutions to reverse the polarity and disrupt hydrophobic interaction. ITC experiments revealed that the GIT1^L271A/279A^ mutant significantly attenuated its interaction with **cmp-17** with the K_d_ value exceeding 3 μM, whereas the GIT1^L271K/L279K^ mutant completely abolished binding, underscoring the importance of these two leucine residues for **cmp-17** engagement (**Figure 5**).

## CONCLUSION

In summary, our study revealed the influence of different ring-closing linkers on the selection outcomes from DNA-encoded cyclic peptide libraries. We constructed eight sub-libraries, each featuring cyclic peptide whose rings were closed using seven bis-electrophilic linkers or disulfide bond. Two targets, MDM2 and GIT1, were screened using theses pooled cyclic peptide libraries, resulting in two distinct yet representative scenarios.

The 3D plots of selection outcomes for MDM2 revealed similar enrichment patterns across four sub-libraries (**DEL-3, DEL-5, DEL-6**, and **DEL-7**), with the BB combination 32_62_390 consistently exhibiting high enrichment values. We selected several high-priority BB combinations, including 32_62_390, for subsequent off-DNA synthesis and validation. Interestingly, compounds derived from the 32_62_390 combination demonstrated good to excellent binding affinities, despite their peptide rings being cyclized by different linkers and exhibiting significantly varied enrichment values across sub-libraries. Drawing definitive conclusions regarding the precise influence of cyclization linkers on selection outcomes remains challenging, as multiple factors—such as DNA tags, PEG spacers, and conformational differences induced by distinct linkers—collectively complicate interpretation. Nevertheless, this finding suggests that structural similarity patterns across multiple hit series or different libraries can enhance confidence in the authenticity of identified hits.

In contrast to MDM2, selection outcomes for GIT1 did not show consistent enrichment patterns across sub-libraries. We identified two BB combinations for further validation through both on-DNA and off-DNA synthesis. These results highlighted another inherent limitation of DEL screening: enriched compounds may not bind to the desired pocket. To address this limitation, parallel selection experiments can be performed when a known positive ligand is available. By adding this ligand to block the desired binding pocket in the selection condition, compounds enriched exclusively in the ligand-free condition can be confidently identified as specific binders. However, if no known positive ligand exists for the target, additional biochemical and biophysical assays are required to validate the authenticity and specificity of selected binders.

Identifying individual hit compounds remains a routine paradigm in DNA-encoded library technology. As researchers gain deeper insights into the field, they increasingly recognize limitations in interpreting DEL selection data, particularly for cyclic peptide entities. The large molecular weights and flexible conformations of cyclic peptides pose additional challenges in accurately identifying authentic binders and their correct binding orientations with protein targets. Integrating computational methods and structural analysis tools represents a promising approach to overcoming these challenges and guiding future discovery efforts.

## ASSOCIATED CONTENT

### Supporting Information

This material is available free of charge via the Internet at Synthetic protocols and MS spectra of DNA-encoded library and on-DNA compounds, selection protocols.

## AUTHOR INFORMATION

**Authors**

**Hanqing Zhao -** *State Key Laboratory of Drug Research, Shanghai Institute of Materia Medica, Chinese Academy of Sciences, Shanghai 201203, P. R. China; University of Chinese Academy of Sciences, Beijing 100049, P. R. China*

**Xue Li -** *Department of Medicinal Chemistry, School of Pharmacy, China Pharmaceutical University, Nanjing 211198, China*

**Xueyu Yao -** *Department of Medicinal Chemistry, School of Pharmacy, China Pharmaceutical University, Nanjing 211198, China*

**Shijie Zhang -** *State Key Laboratory of Drug Research, Shanghai Institute of Materia Medica, Chinese Academy of Sciences, Shanghai 201203, P. R. China; University of Chinese Academy of Sciences, Beijing 100049, P. R. China*

**Yandan Bao -** *State Key Laboratory of Drug Research, Shanghai Institute of Materia Medica, Chinese Academy of Sciences, Shanghai 201203, P. R. China*

**Weiwei Lu -** *State Key Laboratory of Drug Research, Shanghai Institute of Materia Medica, Chinese Academy of Sciences, Shanghai 201203, P. R. China*

**Minyan Xing -** *Department of Medicinal Chemistry, School of Pharmacy, China Pharmaceutical University, Nanjing 211198, China; State Key Laboratory of Drug Research, Shanghai Institute of Materia Medica, Chinese Academy of Sciences, Shanghai 201203, P. R. China*

**Xudong Wang -** *State Key Laboratory of Drug Research, Shanghai Institute of Materia Medica, Chinese Academy of Sciences, Shanghai 201203, P. R. China; University of Chinese Academy of Sciences, Beijing 100049, P. R. China*

**Author Contributions**

^#^ H. Q. and X. L. contributed equally.

**Notes**

The authors declare no competing financial interest.

## ACKNOWLEDGMENT

X.L. was supported by the National Natural Science Foundation of China (NSFC) 22377139, 92253305. Q.C. was supported by National Natural Science Foundation of China (NSFC) 22277141.

